# KilR of *E. coli* Rac prophage is a dual morphogenetic inhibitor of bacterial cell shape

**DOI:** 10.1101/2025.01.07.631774

**Authors:** Anusha Marepalli, Muruganandam Nandhakumar, Sutharsan Govindarajan

## Abstract

Bacterial cryptic prophages encode genes that reduce the viability of the host, upon induction, but also contribute to host survival during stress conditions. Rac is a cryptic prophage of *Escherichia coli* and it encodes a toxic protein KilR which causes morphological defects to the host. But the mechanistic basis of its action is not well understood. In this study, we provide evidence that KilR is a dual morphogenetic inhibitor that affects cell division and cytoskeletal organization. We show that KilR expression is highly toxic, as demonstrated previously, and its predicted C-terminal unstructured region plays a crucial role in its function via a length-dependent manner. Low levels of KilR expression lead to cell filamentation and disruption of Z-rings, while high levels result in rod-shaped defects and mislocalization of the MreB cytoskeletal protein. Using fluorescent fusions, we observed that KilR is diffusively localized in the cytoplasm. When MreBCD proteins are overexpressed, KilR co-localizes with them, forming membrane-associated filaments, indicating a physical association. However, overexpressed MreBCD proteins does not alleviate the KilR-associated growth defect, unlike FtsZ. Finally, we present evidence that chromosomal KilR contributes to the co-inhibition of FtsZ and MreB localization in response to oxidative stress. Our data indicate that KilR inhibits MreB-associated cytoskeletal system, in addition to its effect on FtsZ-associated cell division system. We propose that dual inhibition activity of KilR contributes to its high level of toxicity and to its function in SOS-independent DNA damage tolerance during oxidative stress.

**IMPORTANCE:** KilR is a Rac cryptic prophage encoded toxic protein which contributes to host survival during oxidative stress conditions. It is known to inhibit cell division by targeting the tubulin homolog, FtsZ. In this study, we show that KilR is a dual morphogenetic inhibitor that affects FtsZ-mediated cell division and MreB-mediated cell elongation. Simultaneous inhibition of cell division and cell elongation are known to be crucial for bacterial survival during stress conditions like oxidative stress. Our study identifies KilR as a dual morphogenetic inhibitor, offering insights into how bacterial-phage coevolution drives the emergence of cryptic prophage elements, with specific genes enhancing bacterial fitness.

## INTRODUCTION

Bacterial genomes harbor several cryptic prophages which, unlike prophages, are no longer capable of producing infectious phage particles. These cryptic prophages persist in the bacterial genome as DNA fossil remnants for millions of years and provide selective advantage to the bacterial host. For example, several cryptic prophages encode genes for toxins, virulence factors, or antibiotic resistance and they contribute to the bacterial survival under various stress conditions like oxidative damage, exposure to antibiotics, and during biofilm formation (1, 2).

*Escherichia coli* K-12 harbors nine cryptic prophages, constituting nearly 3.6% of its genome. Rac, the first prophage discovered in *E. coli* (3, 4), is a phage fossil which was acquired over 4.5 million years ago. Rac is a defective lambdoid prophage, measuring 23 kb in length and encompassing 29 genes. It is specifically integrated at the *ttcA* gene, responsible for encoding tRNA-thioltransferase, TtcA. Rac can undergo excision from the *ttcA* locus during normal growth conditions, but this excision is induced under specific circumstances, such as during biofilm formation. Upon Rac excision, an altered TtcA protein (TtcA’) is formed, resulting in a growth defect in the presence of antibiotics like carbenicillin (5, 6). In addition, Rac also contributes to various other phenotypes, including biofilm formation, resistance to nalidixic acid, oxidative stress tolerance, etc (2, 5, 6) highlighting its importance as a cryptic mobile genetic element supporting bacterial survival.

KilR is a toxic protein encoded by the Rac prophage (7, 8). Under normal conditions, expression of most of the genes of the Rac prophage including *kilR* is tightly repressed (9, 10). However, in response to exposure to nalidixic acid or oxidative stress, KilR transcription is transiently induced, which contributes to stress survival. Many of the Rac-associated advantageous phenotypes are attributed to KilR (2, 11).

The role of KilR in bacterial survival during oxidative stress has been well investigated. OxyS sRNA is induced in response to oxidative stress (12). OxyS directly downregulates NusG, a transcription termination factor, through sRNA-mRNA interaction. NusG together with Rho contributes to the repression of KilR. In the presence of OxyS, the reduced expression of NusG, in turn, indirectly leads to the activation of KilR (9). The transiently activated KilR inhibits FtsZ, the major cell division protein, causing temporary growth arrest in an SOS-independent manner. This arrest provides the necessary time for facilitating DNA damage repair and enabling cellular recovery (11). In addition to promoting cell division inhibition through KilR, OxyS was recently shown to inhibit cell elongation through the repression of mepS mRNA, which encodes peptidoglycan endopeptidase (13). Thus, OxyS-mediated co-inhibition of the two interlinked pathway, namely cell division and cell elongation, has been shown to be important for cell survival and recovery from oxidative stress-induced DNA damages.

Several studies have documented that KilR expression leads to cell filamentation, indicating the inhibition of cell division (7, 8, 11). Importantly, increased expression of FtsZ, the central cell division protein, completely suppresses KilR toxicity and associated cell division defects, suggesting an association between KilR and the cell division process (7, 11) Intriguingly, it has also been observed that KilR expression leads to other morphological defects, such as the formation of lemon-shaped cells, which are hallmarks of cell elongation defects (7). Inhibition of FtsZ cannot explain cell elongation defects, indicating that KilR inhibits additional targets that mediate cell elongation. In this study, we provide evidence that KilR is a phage-encoded dual morphogenetic inhibitor, which in addition to inhibiting cell division, can also inhibit cell elongation by targeting the rod shape determining MreBCD cytoskeletal system.

## RESULTS AND DISCUSSION

### KilR mediated toxicity is associated with cell filamentation and rounding

In order to investigate the impact of KilR on *E. coli* growth, we constructed a plasmid expressing KilR under an arabinose-inducible promoter. We assessed the growth of *E. coli* expressing KilR or a control protein, mCherry, by spotting serial dilutions of overnight cultures onto LB agar plates with or without varying concentrations of arabinose. The results, summarized in Fig. 1A, indicate that KilR displayed a concentration-dependent inhibition of cell growth. At a lower arabinose concentration (0.01%), slight defect in cell viability was observed. However, at higher concentrations, growth was completely inhibited, consistent with previous observations of KilR’s toxicity (7). KilR-mediated growth arrest was also evident in liquid growth assays in an inducer concentration-dependent manner (Fig. 1B).

**Fig.1. Growth.**
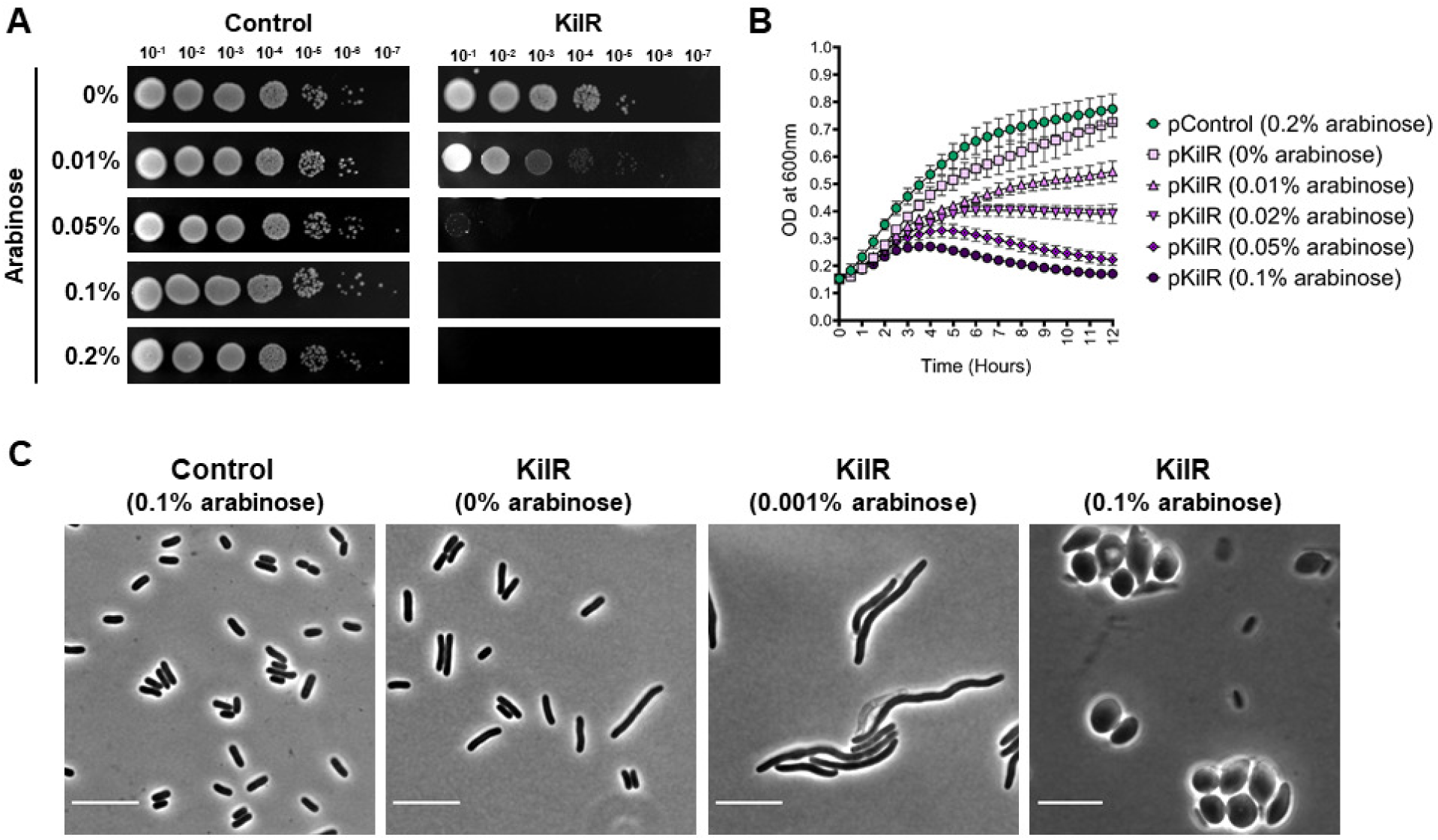
inhibition and morphological effects of KilR expression. (A) Wild-type cells transformed with pBAD-mCherry (control) or pBAD-KilR were spotted in serial dilutions on LB plates without or with the inducer arabinose at varying concentrations and incubated overnight at 37°C. (B) Growth curves of cells expressing KilR in the presence of different concentrations of arabinose or the mCherry control (induced with 0.2% arabinose). (C) Phase contrast images of wild-type cells expressing mCherry control or KilR at various concentrations of inducer arabinose. Scale bar corresponds to 10 μm.

Next, we investigated the morphological effects associated with KilR expression. Expression of KilR at a low concentration of the inducer arabinose resulted in the formation of severely elongated cells, indicating inhibition of cell division. In contrast, in the presence of high concentration of the inducer arabinose, expression of KilR led to the formation of lemon-shaped or spherical cells, a hallmark of cell elongation defects (Fig. 1C, right panel). In the absence of any inducer, cells appeared completely normal (Fig. 1C, left panel). While similar morphological defects were reported for KilR in one study (7), discrepancies exist in other reports in which KilR expression only results in cell filamentation (8, 11). To validate our findings, we assessed morphological defects associated with KilR expression using a previously reported plasmid, pSA97 (11). Consistent with our observations, KilR expressed from pSA97 induced cell filamentation at low concentrations of inducer and formed lemon-shaped or round-shaped cells at high concentrations of inducer (Fig. S1).

The phenotypes associated with KilR overexpression resemble those induced by other proteins targeting both cell division and cell elongation. This includes CptA, a host-encoded toxin part of the CptA-CptB toxin-antitoxin system (14), and CbtA, a toxin encoded in the CP4-44 prophage CbtA-CbeA toxin-antitoxin system (14, 15). Despite lacking homology with KilR, these toxin proteins exhibit comparable morphological effects. Notably, they are part of toxin-antitoxin system and are known to play a crucial role in oxidative stress tolerance (16). However, in contrast, it remains uncertain whether KilR is part of a toxin-antitoxin system. Nevertheless, the results presented above suggest that KilR serves as a dual morphogenetic inhibitor capable of affecting both cell division and cell elongation in *E. coli*.

### Effect of KilR is distinct compared to λ Kil

Similar to the KilR of the Rac prophage, the λ phage also encodes a toxic Kil peptide that affects cell growth (17). Although Rac KilR and λ Kil are not related, both proteins are known to target FtsZ. In addition to FtsZ, λ Kil also inhibits another cell division protein, ZipA (18, 19). FtsA* (R286W) is a gain-of-function mutation which causes an altered FtsA-FtsZ interaction that can bypass the requirement for several cell division proteins including ZipA (20). Interestingly, a previous study has shown that in the mutant background of *ftsA*ΔzipA,* the toxicity of λ Kil is completely abolished (18). Since KilR targets FtsZ similar to λ Kil, we tested whether *ftsA*ΔzipA* background could rescue KilR toxicity. To investigate this, we tested the toxicity of KilR in *ftsA\** and *ftsA*ΔzipA* strains, comparing it with λ Kil. The results presented in Fig. 2A show that the expression of λ Kil in *ftsA\** did not confer resistance to cells. Consistent with a previous report (18), double mutant *ftsA*ΔzipA* cells exhibited complete resistance to λ Kil expression. However, in both *ftsA\** and *ftsA*ΔzipA* strains, KilR expression was toxic to comparable levels (Fig. 2A), suggesting that KilR mediates its function distinctly from λ Kil. To further evaluate this observation, we compared the effects of λ Kil and KilR on *E. coli* cell morphology. In the presence of low concentration of inducer, both λ Kil and KilR caused cell filamentation. However, in the presence of high inducer concentration, cells carrying λ Kil exhibited increased filamentation, whereas cells expressing KilR became round (Fig. 2B). Thus, although Rac KilR and λ Kil target FtsZ, their additional targets are distinct. For λ Kil, ZipA is the additional target, but for KilR the additional target is unidentified. The formation of lemon-shaped or spherical cells is a characteristic feature of cell elongation defects. Such phenotypes are observed in cell elongation-defective mutants disrupted for the MreBCD cytoskeletal system or its associated proteins, including RodZ or RodA (21). Since increased expression of KilR caused these morphological phenotypes, it suggests the possibility of cell elongation-associated cytoskeletal components as additional targets.

**Fig. 2.**
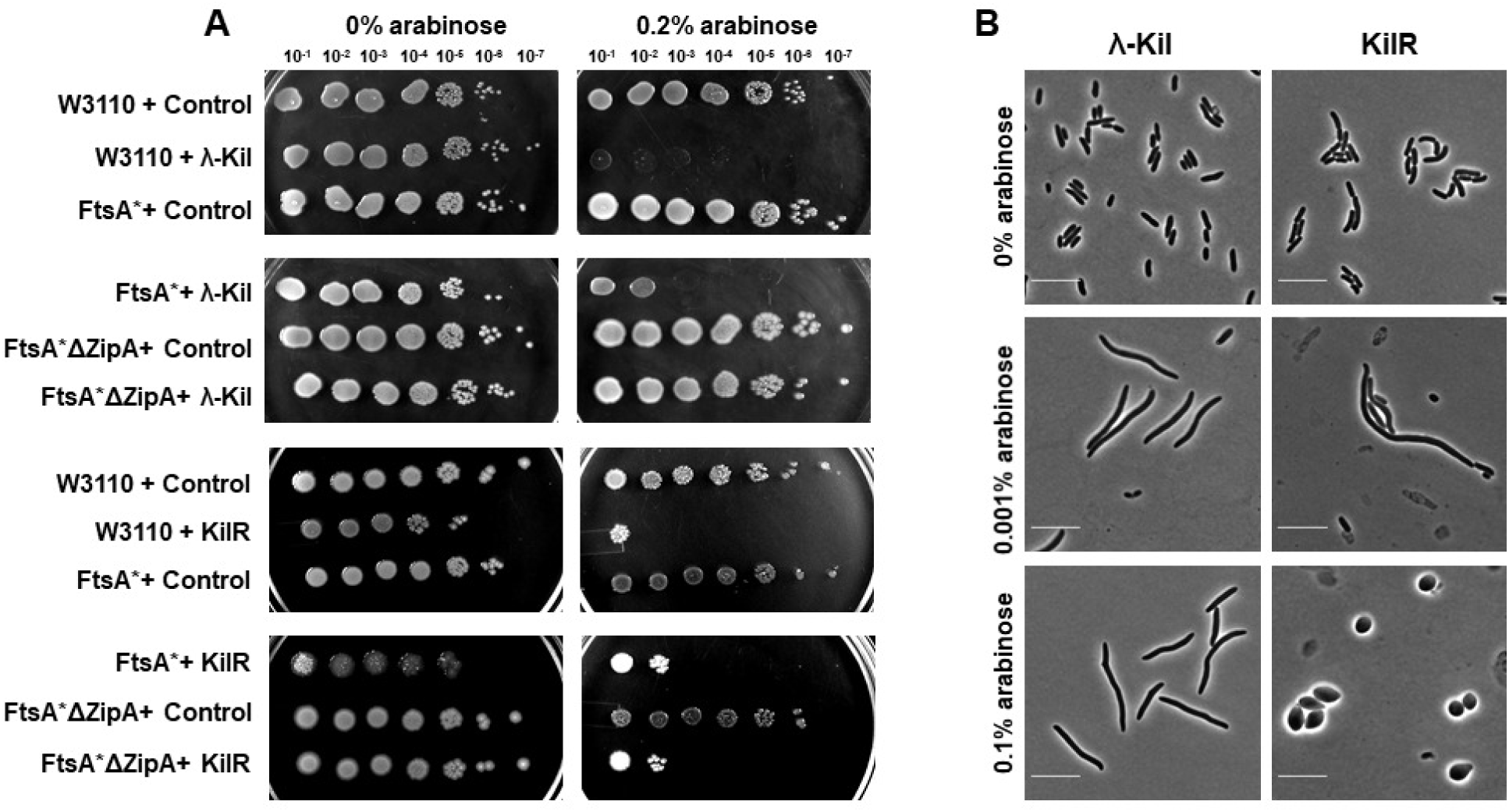
Comparison of λ Kil and Rac KilR. (A) Serial dilutions of the specified strains transformed with plasmids expressing control mCherry, λ Kil, or KilR were cultured under conditions where λ Kil expression is induced (at 42°C) as previously described in (18). Plasmids were either left uninduced or induced with 0.2% arabinose. (B) Phase contrast images of wild-type cells expressing λ Kil (left panel) or KilR (right panel) with different concentrations of inducer arabinose. Scale bar corresponds to 10 μm.

### C-terminal region of KilR is critical for its toxicity and dual morphological effects

To further understand the nature of KilR, we predicted its structure using AlphaFold3 (22). The predicted structure of KilR showed the central region of the protein is largely composed of beta sheets, but two unstructured regions, corresponding to 7 and 12 amino acids, were present in the N-terminal and in the C-terminal, respectively (Fig. 3A). Short unstructured regions at the N-terminus and C-terminus of KilR were also predicted using IUpred (23), a tool for identifying disordered protein regions, using the short disorder option (Fig. S2).

**Fig. 3.**
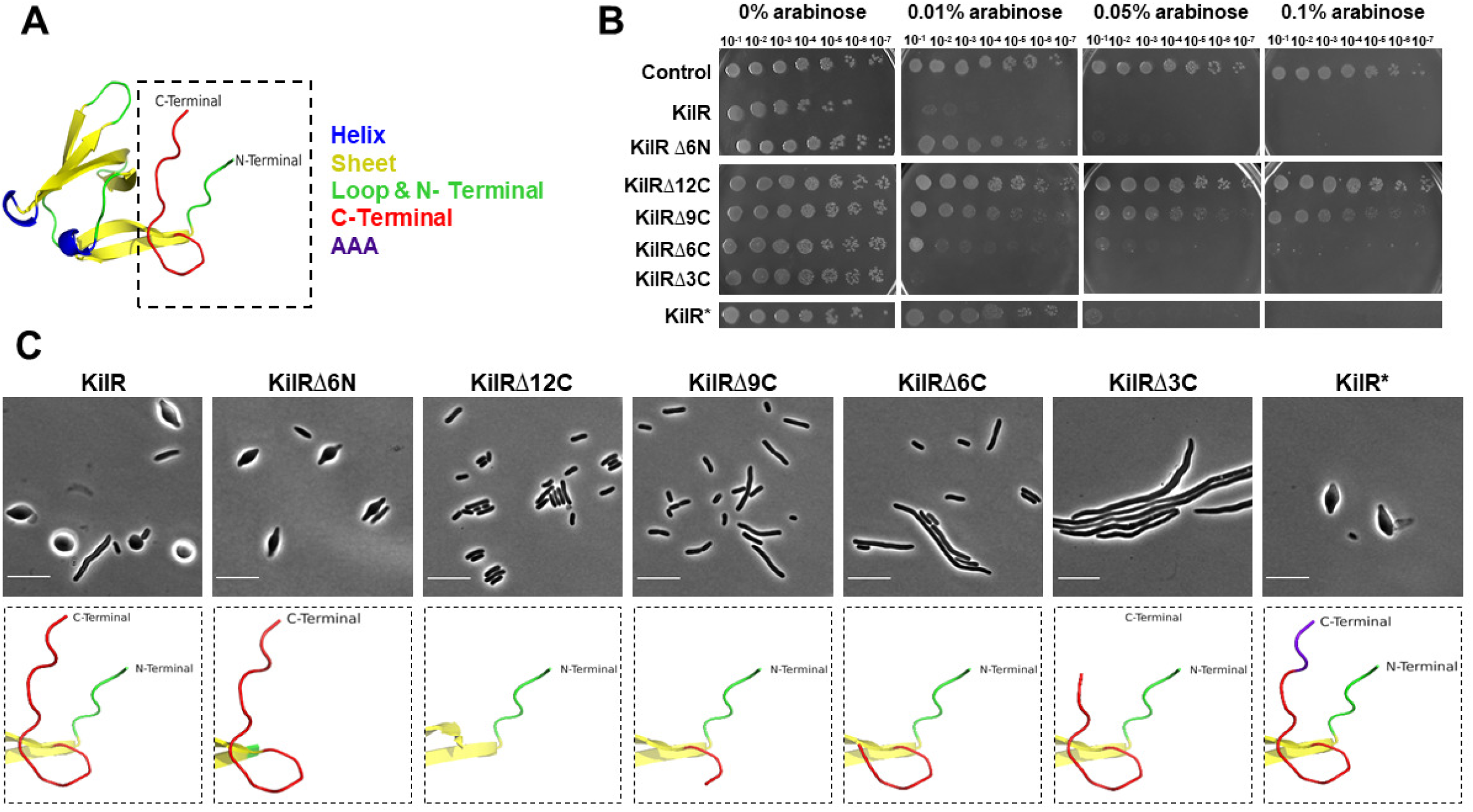
C-terminal unstructured region of KilR is critical for its function. (A) AlphaFold3 predicted structure model of KilR. (B) Serial dilutions of the wild-type transformed with plasmids expressing control mCherry, KilR, KilRΔ6N, KilRΔ12C, KilRΔ9C, KilRΔ6C, KilRΔ3C, and KilR* (3 Ala variant) without or with the inducer arabinose at varying concentrations and incubated overnight at 37°C. (C) Phase contrast images of wild-type cells expressing mCherry control, KilR, or specified KilR variants. Scale bar corresponds to 10 μm. Models depicting the N-terminal and C-terminal regions are shown below the respective KilR versions.

To investigate the importance of these unstructured regions in KilR‘s function, we generated KilR mutants lacking the N-terminal (KilRΔ6N) and the C-terminal (KilRΔ12C) unstructured regions and assessed their impact on toxicity and morphological effects. The results presented in Fig. 3B show that deletion of the N-terminal unstructured region did not affect KilR’s function, as it retained toxicity, similar to the wild-type. In contrast, deletion of the C-terminal unstructured region completely abrogated its toxicity. Morphological analysis supports the toxicity assay, showing that cells expressing KilRΔ6N appeared lemon-shaped, while cells expressing KilRΔ12C appeared completely normal (Fig. 3C).

To further investigate the critical regions within the C-terminal unstructured region, we generated a series of truncations by deleting the last 9 amino acids (KilRΔ9C), 6 amino acids (KilRΔ6C), and 3 amino acids (KilRΔ3C) and assessed their effects. KilRΔ9C and KilRΔ6C showed a gradual reduction in cell viability, with KilRΔ6C still causing slight filamentation (Fig. 3B and 3C). Interestingly, KilRΔ3C, which lacks only the last 3 amino acids, retained its toxicity but lost the ability to induce cell rounding defects, instead leading to the formation of filamented cells (Fig. 3B and 3C). These findings suggest that the last 3 amino acids are essential for inhibiting cell elongation. Western blot analysis of the wild-type and C-terminal truncated proteins, tagged with GST in the N-terminal region, confirmed that all proteins were expressed as full-length proteins, although fragmented proteins were also detected (Fig. S3). These results suggest that the observed differences are not solely due to a lack of protein expression, which can sometimes result from C-terminal mutations (24).

The last 3 amino acids of KilR, namely Glu-Ser-Trp (E-S-W), are largely conserved among homologs of KilR (Fig. S4). We wanted to know whether the specific nature of the last three amino acids is critical for the observed effects. To this end, we created a KilR variant, termed KilR*, in which the last three amino acids were replaced with alanines. Remarkably, KilR* regained the ability to inhibit cell elongation and exhibited toxicity and morphological defects similar to the wild-type KilR (Fig. 3B and 3C).

Collectively, these data support the following conclusions: (a) the C-terminal region of KilR is crucial for mediating its toxicity and represents the active region involved in inhibiting cell division and cell elongation, (b) the gradual reduction of toxicity observed with progressive truncation of the C-terminal unstructured region suggests that its length contributes to the strength of KilR toxicity, and (c) the last 3 amino acids, irrespective of their sequence, potentiate KilR’s ability to inhibit cell elongation, thereby enabling its dual-targeting nature. Additionally, deletion of the last 3 amino acids allowed us to uncouple the cell division and cell elongation defects caused by KilR without compromising its toxicity. Proteins with C-terminal minimotifs have been proposed to mediate additional molecular recognition (25), and our results add KilR to this category. However, mechanistic insights will require detailed interaction and structural analysis of KilR and its targets.

### KilR perturbs Z-ring formation and MreB localization

Having observed that KilR expression causes cell division as well as cell elongation defects, we next checked the effect of KilR and its truncated variants on the sub-cellular organization of FtsZ and MreB, which mediate cell division and cell elongation, respectively (26–29). To answer this question, we used a strain that expressed ZapA-GFP and MreB-RFP^SW^ from the native locus (30–32). ZapA is an early cell division protein which co-localizes with FtsZ and can be reliably used as a marker for FtsZ localization (31, 33, 34).

The results, presented in Fig. 4, reveal that KilR overexpression caused significant perturbations in the localization patterns of both ZapA-GFP and MreB-RFP^SW^. Specifically, ZapA-GFP exhibited a spotty appearance, losing its characteristic Z-ring formation, while MreB-RFP^SW^ formed aberrant, mislocalized clusters. In contrast, control cells displayed clear mid-cell localization of ZapA-GFP and membrane-associated spotty localization of MreB-RFP^SW^, as previously observed (32).

**Fig. 4.**
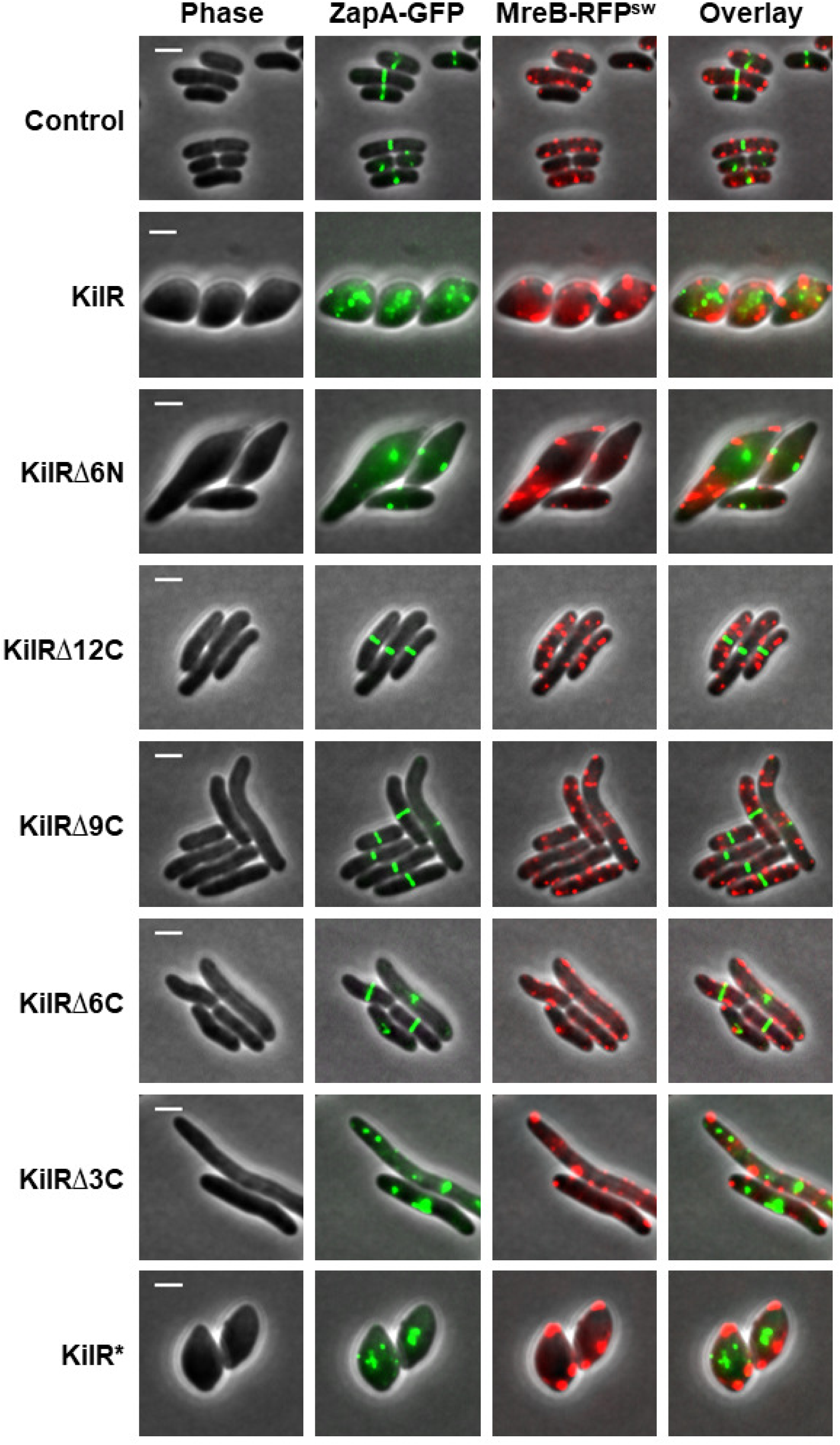
KilR overexpression affects the localization of FtsZ and MreB. Fluorescence microscopy images of cells expressing Zap-GFP and MreB-RFP^SW^ and transformed with plasmids expressing control (GST), wild-type KilR, its truncated versions (KilRΔ6N, KilRΔ12C, KilRΔ9C, KilRΔ6C, or KilRΔ3C), or KilR* (3 Ala variant). The GFP and mCherry fusion proteins were observed by fluorescence microscopy and cells were observed with phase contrast microscopy. Shown are phase contrast (grey), GFP (green), and mCherry (red) fluorescence signals. Fluorescent signals are overlaid on phase contrast images. Scale bar corresponds to 2 μm

Cells expressing KilRΔ6N showed disruptions in ZapA-GFP and MreB-RFP^SW^ localization, similar to wild-type KilR. Conversely, in cells expressing KilRΔ12C, both ZapA-GFP and MreB-RFP^SW^ localizations appeared completely normal, indicating that the absence of the last 12 amino acids abolished KilR’s inhibitory effects on Z-ring formation and MreB localization. Cells expressing KilRΔ9C and KilRΔ6C exhibited partial effects: ZapA-GFP formed Z-rings in most cells, except for those displaying slight elongation, and MreB-RFP^SW^ localization appeared largely normal. In contrast, expression of KilRΔ3C significantly disrupted ZapA-GFP localization but resulted in a mixed pattern of MreB-RFP^SW^ localization, with aberrant clusters alongside normal membrane-associated small spotty clusters. This observation explains the maintenance of rod-shaped morphology during KilRΔ3C overexpression. Finally, cells expressing KilR* disrupted both ZapA-GFP and MreB-RFP^SW^ localization patterns, mirroring the effects of wild-type KilR (Fig. 4). Unlike KilR, the expression of λ Kil induces cell filamentation without disrupting MreB localization (Fig. S5). Taken together, these findings suggest that KilR, via distinct regions of its C-terminal region, modulates the subcellular organization of FtsZ and MreB, thereby influencing cell division and cell elongation, respectively.

### Overexpression of MreBCD recruits KilR but fails to counteract KilR-mediated cell death

To investigate the cellular distribution of KilR and its interactions with cytoskeletal components, we engineered a fluorescently tagged version of KilR by fusing mCherry to its N-terminus. The mCherry-KilR fusion protein retained its toxic effects (Fig. S6) and caused morphological defects similar to untagged KilR (Fig. 5A), indicating that the fluorescent tag did not impair KilR function. Fluorescence microscopy showed that mCherry-KilR exhibited diffuse cytoplasmic localization, similar to the control protein mCherry (Fig. 5A). Western blot analysis confirmed that mCherry-KilR was expressed as a single fusion protein, further supporting that the observed diffuse localization reflected the distribution of KilR in cells. Furthermore, western blot analysis clearly indicates that KilR is expressed in lower levels under conditions of cell filamentation, and at relatively higher levels during cell rounding (Fig. 5B).

**Fig. 5.**
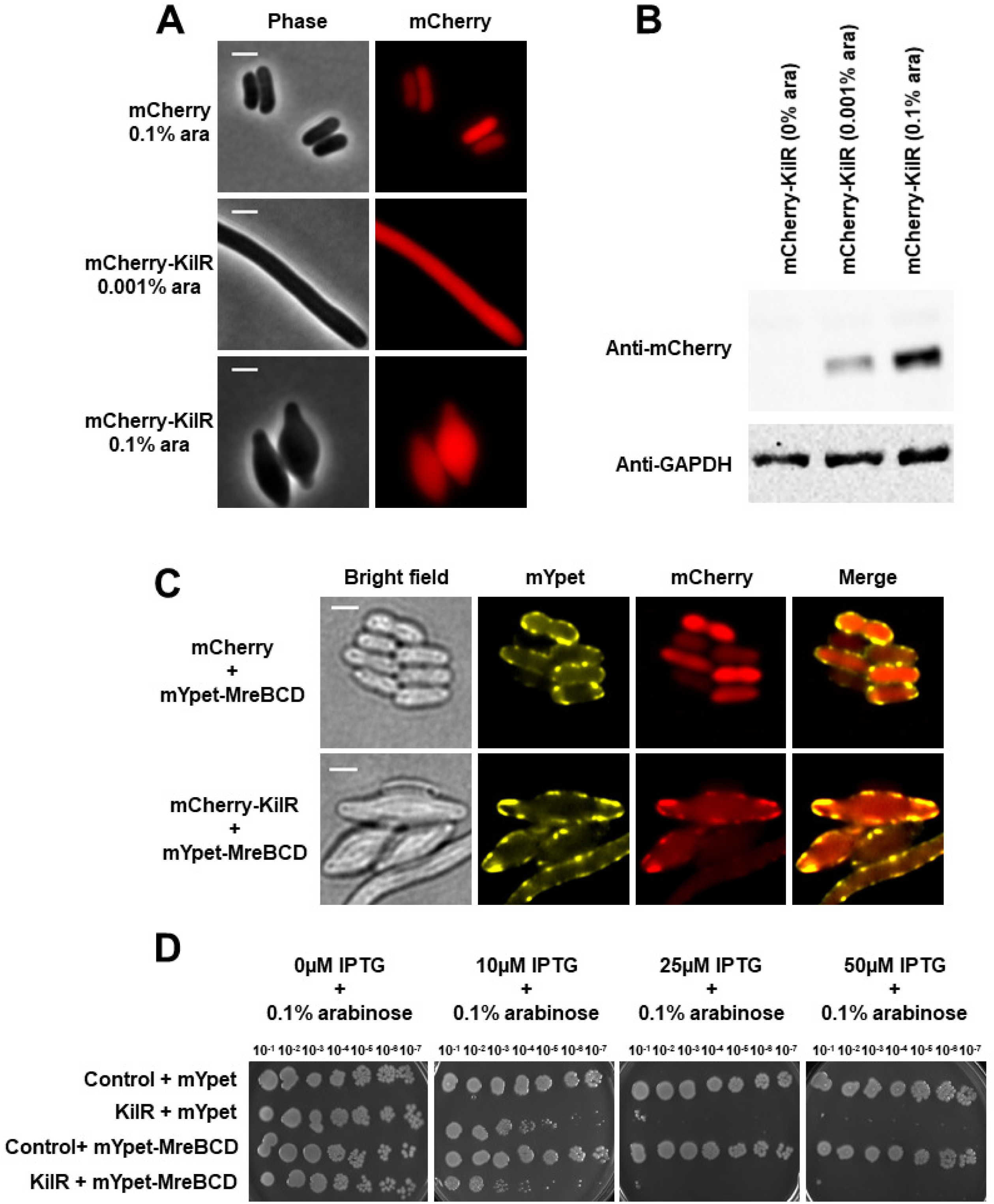
Effect of MreBCD overexpression on KilR localization and function. (A) Fluorescence microscopy images of wild-type cells expressing mCherry, or mCherry-KilR with indicated arabinose concentration. The mCherry was observed by fluorescence microscopy and cells were observed with phase contrast imaging. Shown are phase contrast (grey), and mCherry (red) fluorescence signal. (B) Western blot analysis shows the expression of the mCherry-KilR uninduced or induced with specified concentrations of arabinose (top panel). Western blotting against GAPDH was performed as the housekeeping control (lower panel). (C) Fluorescence microscopy images of wild-type cells containing mYpet-MreBCD and co-expressing mCherry or mCherry-KilR, and induced with 0.4% arabinose. The mYpet and mCherry fusion proteins were observed by fluorescence microscopy and cells were observed with phase contrast microscopy. Shown are phase contrast (grey), and mYpet (yellow), and mCherry (red) fluorescence signals. Scale bar corresponds to 2 μm. (B) Serial dilutions of the wild-type transformed with specified plasmids and grown without or with the inducer arabinose at varying concentrations and incubated overnight at 37°C.

The MreBCD proteins form a membrane-associated complex that co-localizes with interacting proteins such as RodZ, MbiA, etc (35–37). We hypothesized that KilR might associate with the MreBCD complex, influencing its subcellular distribution. To test this, we co-expressed mYpet-tagged MreBCD along with either mCherry-KilR or mCherry alone and analyzed their localization patterns. As expected, mYpet-MreBCD formed membrane-associated filaments when overexpressed from a plasmid, consistent with previous reports (38). Remarkably, in cells co-expressing mCherry-KilR and mYpet-MreBCD, mCherry-KilR co-localized with the membrane-associated MreBCD filaments (Fig. 5C. lower panel). In contrast, control cells expressing mCherry alone showed no co-localization with mYpet-MreBCD (Fig. 5C, upper panel). This observation indicates that KilR associates with the MreBCD complex.

Next, we assessed whether MreBCD overexpression could counteract the toxic effects of KilR. As shown in Fig. 5D and Fig. S7, overexpressing MreBCD did not mitigate KilR-mediated toxicity. This contrasts with the complete suppression of KilR toxicity and associated morphological defects during FtsZ overexpression (7, 11). Our findings suggest that KilR associates with the MreBCD cytoskeletal complex, but the increased concentration of MreBCD proteins fails to inhibit KilR toxicity. This could be attributed to reduced affinity of KilR and MreBCD, causing ineffective sequestration, or the continued inhibition of FtsZ even in the presence of high concentrations of MreBCD proteins, or a combination of both.

### KilR contributes to dual inhibition of FtsZ and MreB during oxidative stress

Under normal growth conditions, *kilR* is tightly repressed (9), but its expression is induced during specific stress conditions such as oxidative stress (11). Since plasmid-expressed KilR can inhibit both processes, our next investigation focused on whether a similar effect could be observed for the native chromosomal KilR, through induction of oxidative stress. To explore this, we examined the localization of FtsZ-mCherry, expressed from a plasmid, and MreB-msfGFP^SW^, expressed from the native locus, in both wild-type and *ΔkilR* mutant strains. Oxidative stress was induced by adding various concentrations of H_2_O_2_, and the effects on protein localization were assessed.

In wild-type cells, oxidative stress caused a clear disruption of both FtsZ-mCherry and MreB-msfGFP^SW^ localization compared to untreated cells. Mislocalization of FtsZ-mCherry was evident at even low concentrations of H₂O₂ (3 mM), whereas MreB mislocalization occurred only at higher concentrations (5 mM and 7 mM) (Fig. 6A).

**Fig. 6.**
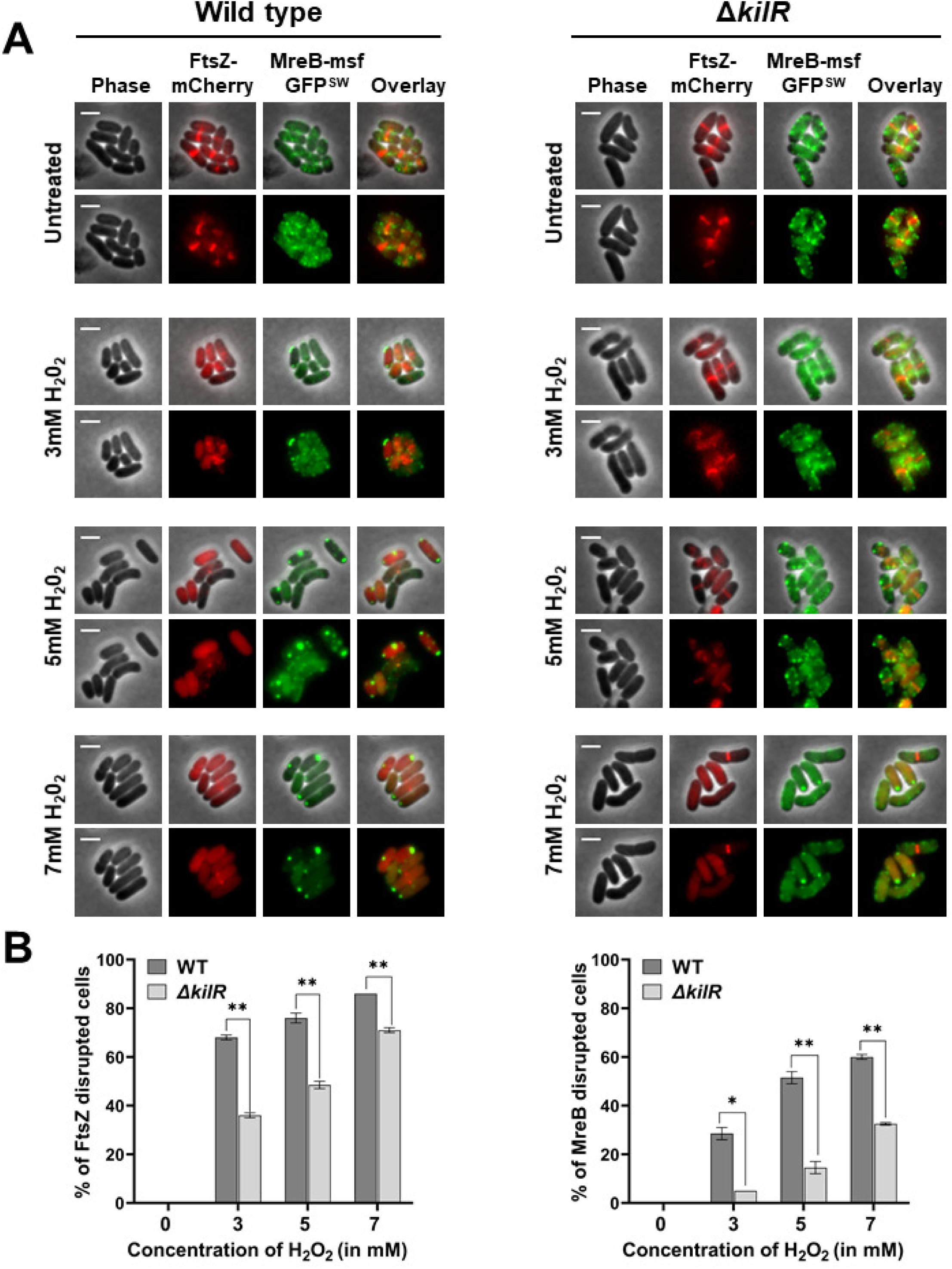
KilR affects FtsZ and MreB localization during oxidative stress. (A) Fluorescence microscopy images of wild-type and *ΔkilR* cells expressing FtsZ-mCherry and MreB-msfGFP^SW^ and untreated or treated with H_2_O_2_ at various concentrations. The mCherry and GFP fusion proteins were observed by fluorescence microscopy and cells were observed with phase contrast microscopy. Shown are phase contrast (grey), and mCherry (red), and GFP (green) fluorescence images as well as overlay over phase contrast images. Scale bar corresponds to 2 μm. (B) Percentage of cells with mislocalized FtsZ-mCherry (left) or MreB-msfGFP^SW^ (right) are shown. The data represents approximately 200 cells analyzed from two independent experiments. Means and standard deviations are shown. The statistical significance was calculated using unpaired t-test analysis (* p<0.05, ** p<0.01).

Interestingly, in the Δ*kilR* background, the mislocalization of both FtsZ-mCherry and MreB-msfGFP^SW^ was less pronounced following H₂O₂ treatment (Fig. 6A and 6B). For FtsZ-mCherry, significant differences were observed at all concentrations of H₂O₂, whereas for MreB-msfGFP^SW^, differences were more significant at 5 mM and 7 mM H₂O₂ concentrations (Fig. 6B). Overall, these findings suggest that KilR, when expressed under physiologically relevant stress conditions such as oxidative stress, mediates the dual inhibition and mislocalization of FtsZ and MreB. The observed effect is stronger for FtsZ compared to MreB. Furthermore, under the tested conditions, we observed disruptions in the localization of FtsZ and MreB without inducing any morphological defects.

## CONCLUSION

The results of this study establish KilR as a phage-encoded morphogenetic inhibitor capable of dual inhibition of FtsZ-mediated cell division and MreB-mediated cytoskeletal processes. Characterization of KilR reveals that a 12-amino acid unstructured region in the C-terminus, as predicted by AlphaFold3, to be important for its dual inhibitory function. However, a KilR variant lacking the last 3 amino acids lost its dual-targeting ability, resulting in inhibition of only the cell division process. Replacing these 3 amino acids with alanines restored KilR’s dual-targeting activity. These observations suggest that the length of the unstructured region of the C-terminus provides flexibility, enabling interactions with multiple targets. Localization studies demonstrated that KilR diffuses cytoplasmically but associates with the MreBCD complex upon its overexpression. Despite this co-localization, elevated MreBCD levels failed to counteract KilR-mediated cell death, suggesting a dominant inhibitory effect of KilR, presumably due to its dual inhibitory effect on FtsZ-associated cell division and MreB-associated cytoskeletal processes.

Dual targeting of cell division and cytoskeletal organization has recently been proposed to contribute to bacterial survival and their ability to cope with DNA damage during oxidative stress (13). The OxyS small RNA, induced in response to oxidative stress, possesses antimutator properties by reducing bacterial DNA damage capacity after exposure to oxidative stress (11, 12). Previous studies have indicated that dual inhibition of cell division and the cytoskeleton is orchestrated by two distinct pathways of OxyS. In one pathway, OxyS induces KilR, which causes SOS-independent blockade of cell division (11), while in another pathway, OxyS reduces the expression of *mepS*, the gene encoding a peptidoglycan endopeptidase crucial for cell elongation (13). Our findings, revealing that KilR itself is capable of dual targeting, suggest the redundancy of this pathway and underscore its significance in oxidative stress tolerance.

## MATERIALS AND METHODS

### Bacterial strains and growth media

Strains and plasmids used in this study are listed in supplementary Table S1. Overnight *E*. *coli* cultures were grown in LB at 37°C or 30°C, depending on the experiment, supplemented with appropriate antibiotics. When appropriate, antibiotics were added at the following concentrations: ampicillin (100 μg/ml), kanamycin (50 μg/ml), or chloramphenicol (30 μg/ml). Antibiotics, arabinose, and IPTG were purchased from Sisco Research Laboratories.

### Construction of strains and plasmids

AM001, which expresses MreB-msfGFP^SW^ in *ΔkilR* background, was constructed by P1-transduction transferring mreB’-msfGfp-‘mreBcsrD(::neo) from NO50 (39) to A870 (11). Plasmids expressing mCherry or KilR were constructed by Gibson assembly in the pBAD18 plasmid. mCherry was amplified from pSG30T-sfCherry-csy1(IF) (40) using the primers F-pBAD18-rbs-cherry and R-pBAD18-mcherry, and kilR was amplified from the genomic DNA of *E. coli* MG1655 K12 using F-vec-rbs-Kil and R-vec-kilR primers. The resulting amplicons were Gibson assembled in the pBAD18 vector backbone which was amplified using F-pBAD18-gib and R-pBAD18 gib primers. C-terminal truncations of KilR were made as follows: pBAD-KilRΔ3C was made by Gibson ligation using kilR amplified with F-vec-rbs-Kil and R-KilRΔ3C primers, and the pBAD18 vector backbone was amplified using F-pBAD18-gib and R-pBAD18 gib primers. pBAD-KilRΔ6C, pBAD-KilRΔ9C, and pBAD-KilRΔ12C were constructed in a similar manner. However, in all these plasmids, the forward primers for the kilR gene was the same (F-vec-rbs-Kil), and the reverse primers were R-KilRΔ6C (for pBAD-KilR Δ6C), R-KilRΔ9C (for pBAD-KilR Δ9C), and R-KilRΔ12C (for pBAD-KilR Δ12C). N-terminal truncation of KilR and KilR* were made as follows: kilR was amplified using F-KilRΔ6N, R-vec-kilR and F-vec-rbs-Kil, R_KilR* primers respectively. The resulting amplicons were Gibson assembled in the pBAD18 vector backbone which was amplified using F-pBAD18-gib and R-pBAD18 gib primers. pBAD-GST plasmid was constructed by amplifying GST from pGex-6p-2 using F-pBAD18-GST and R-pBAD18-gst, amplicons were assembled into pBAD18 vector i.e., amplified using F-pBAD18-gib and R-pBAD18 gib primers.

GST KilR fusions were constructed as follows: For pBAD-GST-KilR, GST was amplified along with vector using F-pBAD18-gib and R-Lin-GST primers. KilR was amplified without a start codon using F-GST-KilR (-ATG) and R-vec-kilR primers. pBAD-GST-KilRΔ3C was made by Gibson ligation using kilR amplified with F-GST-KilR and R-KilRΔ3C primers, and the pBAD18 vector backbone was amplified along with GST using F-pBAD18-gib and R-Lin-GST primers. pBAD-GST-KilRΔ6C, pBAD-GST-KilRΔ9C, and pBAD-GST-KilRΔ12C were constructed in a similar manner. However, in all these plasmids, the forward primers for the kilR gene was the same (F-GST-KilR), and the reverse primers were R-KilRΔ6C (for pBAD-KilR Δ6C), R-KilRΔ9C (for pBAD-KilR Δ9C), and R-KilRΔ12C (for pBAD-KilR Δ12C).

pBAD-mCherry-KilR was constructed as follows: For pBAD mCherry-KilR, mCherry was amplified without the stop codon using F-pBAD18-rbs-cherry, and R-KilR-SfCherry (-TAA) primers. KilR was amplified without start codon using LIN-KilR Gib (-ATG) and R-vec-kilR primers. mCherry-KilR, GST-KilR and C-terminal truncations of KilR with GST fusion have a ggaggcggtggagcc (G-G-G-G-A) linker sequence between them.

### Spot titers and Growth curve assays

For spot titer assays, overnight cultures were serially diluted 10-fold in fresh LB medium. Subsequently, 3µl of the serially diluted culture was spotted onto LB plates containing the appropriate antibiotic(s) and inducers. The plates were then incubated at the specified temperatures, and images were captured using the Gel Documentation System from Bio-Rad. For the growth curve analysis, overnight cultures were diluted to 1:1000 and grown for 2 hours in fresh LB at 37°C with the appropriate antibiotic. After 2 hours, the inducer arabinose was added at specified concentrations. Then, 180 µl from each sample was aliquoted into a fresh 96-well plate and sealed with Microplate Seal Film, which was punctured with a sterile needle for aeration. Subsequently, the 96-well plate was loaded into a Spark® multimode microplate reader, and optical density (OD) was measured at 600 nm every 15 minutes for a duration of 12 hours. The resulting graph was plotted using GraphPad Prism.

### Expression analysis of GST and mCherry-tagged proteins using western blotting

HB101 *E. coli* cells were transformed with plasmids expressing GST-KilR and C-terminal truncations of KilR fused with GST and purified using GST Sepharose beads according to the manufacturer’s instructions (Cytiva). GST-tagged proteins were expressed as previously described (41). Overnight cultures were diluted to 1:50 in LB media containing ampicillin and grown for 2 hours. After 2 hours, cells were induced with 0.05% arabinose for 4 hours. Cells were harvested by centrifugation at 5000 rpm for 5 minutes. Pellets were stored at −20°C until further use. Pellets were resuspended with lysis buffer pH 7.4 (1X PBS + 1mM PMSF) and sonicated on ice with 30% amplitudes (pulse on 2 seconds and pulse off 5 seconds) till the supernatant was clear, and the supernatant was cleared by centrifugation at 5000rpm/10min. Supernatant was incubated with GST beads for 90 minutes at room temperature (<20°C), followed by three washes with lysis buffer + 300mM NaCl at 2100 rpm for 5 minutes. After thorough washing, the beads were resuspended in 1X SDS sample buffer and subjected to SDS-PAGE.

Overnight cultures of *E. coli* cells expressing mCherry-KilR were diluted 1:5 in LB media containing ampicillin and induced with 0%, 0.001%, and 0.1% arabinose for 4 hours. After induction, equal volumes of samples were collected, washed, and prepared for analysis. Protein samples were separated on a 12% SDS-PAGE gel and transferred to a 0.45 μm nitrocellulose membrane (Cytiva) using the Bio-Rad Mini-PROTEAN Tetra Vertical Electrophoresis system in 1X Tris/Glycine Buffer. The membranes were blocked with 5% BSA in TBST and then probed with primary antibodies diluted 1:10,000 in 1X PBS with 1.25% BSA: Mouse anti-GST (Invitrogen, Cat #MA4-004), Rat anti-mCherry (Invitrogen, Cat #MA5-32977), and Mouse anti-GAPDH (Invitrogen, Cat #MA5-15738). HRP-conjugated Goat anti-Mouse IgG (Invitrogen, Cat #A28177) was used as the secondary antibody. Blots were developed using ECL Start Western Blotting Detection Reagent (Cytiva), and chemiluminescence was detected using a Bio-Rad gel documentation system.

### Structure prediction and multiple sequence alignment

The amino acid sequence of KilR from *E. coli* K12 MG1655 was retrieved from UniProtKB (Accession number: P38393). Alphafold3 was used to predict the KilR protein structure using default parameters (https://alphafoldserver.com/fold/6fa1f4f62ae1e06b) (22). The resulting 3D structure was visualized and analyzed with PyMol (The PyMOL Molecular Graphics System, Version 3.0 Schrödinger, LLC). Truncated protein structures were generated using PyMOL to explore the structural variations. Additionally, the unstructured regions of KilR were predicted using IUPred2A (https://iupred2a.elte.hu/), employing the IUPred2 short disorder option (23).

For Multiple Sequence Alignment analysis, a BLASTP search was conducted against the NR database (*Escherichia coli*) to identify homologs of the Rac KilR protein. This resulted in 842 hits. These hits were analyzed to assess the level of amino acid conservation. Sequence alignment was performed using MUSCLE v3.8.1551 (42), and model selection was carried out using MEGAX v11.0.13 (43). Maximum likelihood phylogenies of the aligned sequences were generated using RAxML v8.2.12 (44) with the PROTGAMMAI (JTT+G4) amino acid substitution model (−m) and 100 bootstrap replicates. The resulting phylogenetic tree was visualized and annotated using the online platform iTOL (45).

### Fluorescence microscopy

Fluorescence microscopy was carried out as described previously (32). Following conditions were utilized to visualize morphological changes upon expression of KilR or its truncated variants: Overnight cultures containing appropriate plasmids were grown in LB at 30°C and supplemented with appropriate antibiotics, then sub-cultured at 1:5 dilutions in fresh LB medium. Cultures were induced with the specified concentration of arabinose and continued to grow for 4 hours with slow agitation (100 rpm). Both induced and uninduced samples were collected after 4 hours and prepared for microscopy.

To study KilR localization using pBAD-mCherry-KilR, cells were grown in LB at 30°C and supplemented with the appropriate antibiotic. The cells were sub cultured at a 1:5 dilutions in fresh LB medium. Cultures were induced with 0%, 0.001%, 0.1% arabinose and continued to grow with slow agitation for 4 hours. Samples were collected after 4 hours and prepared for microscopy.

For visualizing MreB-RFP^SW^ and ZapA-GFP in the presence of KilR and its variants, strain SUT106 expressing both proteins from the native chromosomal loci was transformed with appropriate plasmids. Overnight cultures were grown in LB at 30°C and supplemented with appropriate antibiotics, then sub-cultured at a 1:5 dilutions in fresh LB medium and continued to grow at 30°C with 0.1% arabinose for 4 hours with slow agitation.

For visualizing mCherry or mCherry-KilR together with mYpet-MreBCD, cells containing specified plasmids were grown in LB at 37°C and supplemented with the appropriate antibiotic. The overnight cultures were sub cultured at a 1:50 dilution in fresh LB medium and grown at 30°C for 2 hours with shaking. Cultures were induced with 0.4% arabinose and continued to grow with slow agitation for 3 hours. Samples were collected and prepared for microscopy

To image the localization of MreB and FtsZ under oxidative stress, NO50 and its *ΔkilR* derivative, both containing MreB-msfGFP^SW^ from the native chromosome loci, was transformed with the FtsZ-mCherry plasmid. Overnight cultures were grown in LB at 37°C and supplemented with appropriate antibiotics. Cells were subcultured at a 1:50 dilution in fresh LB medium, then allowed to grow at 30°C for 2 hours. H_2_O_2_ concentrations of 0 mM, 3 mM, 5 mM, and 7 mM were added for 1 hour and cells were allowed to grow at 30°C without agitation. Cells with mis-localized FtsZ and mis-localized MreB were identified from each field of view manually with the support of NIS Elements software. Mislocalized FtsZ refers to cells in which Z-ring is absent and FtsZ-mCherry signal appeared diffuse. Mislocalized MreB refers to cells with aberrant clusters which are mostly mislocalized to the cell poles and cells with diffuse MreB-msfGFP^SW^ signal. Final data were analyzed and represented as a bar graph using GraphPad Prism. All samples for microscopy were washed with 1X PBS. All processed samples were spotted on 0.8% agarose pads prepared with 1X PBS and visualized and photographed using a Nikon Eclipse Ti2-E equipped with a 100X CFI Plan Apochromat Oil objective and a DSQi-2 Monochrome camera.

## Acknowledgment

Sutharsan Govindarajan acknowledges support from DST-SERB (CRG/2020/003295), DBT-Wellcome Trust Early Career Fellowship (IA/E/19/1/504958) and support provided by SRM University – AP (SRMAP/URG/E&PP/2022-23/018 and SRMAP/URG/GENERAL/2024-25/039). Anusha Marepalli and Muruganandam Nandhakumar acknowledge the PhD Research Fellowship from SRM University - AP. We thank Shoshy Altuvia (Hebrew University of Jerusalem) for critical reading of the manuscript. We acknowledge the following individuals for gifts of strains: Durga Rao (SRM University – AP), Orna Amster-Choder and Shoshy Altuvia (Hebrew University of Jerusalem), William Margolin (University of Texas McGovern Medical School). We thank Sharayu Magar for computational analysis and we appreciate helpful discussions with members of the Department of Biological Sciences, SRM University – AP.

## Views

The views expressed in this study are those of the authors and not necessarily those of either the funding agency or any other institution.

## Author contributions

S.G, A.M and M.N formulated the study design and plans. A.M and M.N performed the experiment. S.G wrote the manuscript.

## Supplementary Figure Legends

Fig. S1. Assessing the morphological defects of KilR expressed from pSA97 Images of cells expressing KilR from pSA97 were induced with 0, 5μM and 1mM of IPTG. cells were observed with phase contrast (grey) microscopy.

Fig. S2. Predicting of disorder regions of KilR using IUpred2a IUPred2a prediction of KilR (Unitprot ID - P38393) using the “IUPred2a short disorder” option. Disorder is predicted at the C-terminal and N-terminal regions of the protein.

Fig. S3. Western blot analysis of GST-tagged wild-type KilR and C-terminal truncations of KilR Western blot analysis shows the expression of the control protein (GST), GST-tagged KilR and its C-terminal trucnations. Western blotting against GAPDH was performed as the housekeeping control (lower panel).

Fig. S4. Phylogenetic tree and multiple sequence alignment of KilR and its homologs in *E. coli* Maximum likelihood phylogenetic tree depicting the relationships among KilR protein homologs identified from the BLASTP search. Branch lengths are proportional to the evolutionary divergence, and bootstrap values are displayed to indicate node support. This tree highlights the conservation of the KilR protein across homologs within the *Escherichia coli* dataset. Query sequence is highlighted in red in the phylogenetic tree. Red asterisk (***) are shown in the bottom to indicate the last 3 amino acids (E-S-W) of Rac KilR.

Fig. S5. MreB localization is not affected by λ-Kil Images of MreB-msfGFP^SW^ cells expressing λ-Kil or KilR with or without inducer arabinose. The GFP fusion protein was observed by fluorescence microscopy and cells were observed with phase contrast microscopy.

Fig. S6. mCherry tagging at the N-terminus does affect the toxicity of KilR Cells expressing control (mCherry), KilR, and mCherry-KilR were spotted after serial dilutions on LB plates containing ampicillin with or without arabinose and grown for overnight at 37°C.

Fig. S7. Overexpression of MreBCD does not alleviate KilR toxicity Growth curve of cells expressing KilR or control with mYpet or mYpet-MreBCD. KilR and control are induced with different concentrations of IPTG (0, 10, 50μM), mYpet and mYpet-MreBCD were induced with 0.2% arabinose.

## Supplementary Table

Table S1 – List of strains used in this study

Table S2 – List of plasmids used in this study

Table S3 – List of primers used in this study

## REFERENCES

1. Casjens S. 2003. Prophages and bacterial genomics: what have we learned so far? Mol Microbiol 49:277–300.

2. Wang X, Kim Y, Ma Q, Hong SH, Pokusaeva K, Sturino JM, Wood TK. 2010. Cryptic prophages help bacteria cope with adverse environments. Nat Commun 1:147.

3. Kaiser K, Murray NE. 1979. Physical characterisation of the “Rac prophage” in E. coli K12. Mol Gen Genet MGG 175:159–174.

4. Low B. 1973. Restoration by the rac locus of recombinant forming ability in recB− and recC− merozygotes of Escherichia coli K-12. Mol Gen Genet MGG 122:119–130.

5. Hong SH, Wang X, Wood TK. 2010. Controlling biofilm formation, prophage excision and cell death by rewiring global regulator H-NS of Escherichia coli. Microb Biotechnol 3:344–356.

6. Liu X, Li Y, Guo Y, Zeng Z, Li B, Wood TK, Cai X, Wang X. 2015. Physiological function of rac prophage during biofilm formation and regulation of rac excision in Escherichia coli K-12. Sci Rep 5:16074.

7. Conter A, Bouche J-P, Dassain M. 1996. Identification of a new inhibitor of essential division gene ftsZ as the kil gene of defective prophage Rac. J Bacteriol 178:5100– 5104.

8. Burke C, Liu M, Britton W, Triccas JA, Thomas T, Smith AL, Allen S, Salomon R, Harry E. 2013. Harnessing single cell sorting to identify cell division genes and regulators in bacteria. PLoS One 8:e60964.

9. Cardinale CJ, Washburn RS, Tadigotla VR, Brown LM, Gottesman ME, Nudler E. 2008. Termination factor Rho and its cofactors NusA and NusG silence foreign DNA in E. coli. Science (80-) 320:935–938.

10. Krishnamurthi R, Ghosh S, Khedkar S, Seshasayee ASN. 2017. Repression of YdaS toxin is mediated by transcriptional repressor RacR in the cryptic rac prophage of Escherichia coli K-12. Msphere 2:10–1128.

11. Barshishat S, Elgrably-Weiss M, Edelstein J, Georg J, Govindarajan S, Haviv M, Wright PR, Hess WR, Altuvia S. 2018. OxyS small RNA induces cell cycle arrest to allow DNA damage repair. EMBO J 37:413–426.

12. Altuvia S, Weinstein-Fischer D, Zhang A, Postow L, Storz G. 1997. A small, stable RNA induced by oxidative stress: role as a pleiotropic regulator and antimutator. Cell 90:43–53.

13. Elgrably-Weiss M, Hussain F, Georg J, Shraiteh B, Altuvia S. 2024. Balanced cell division is secured by two different regulatory sites in OxyS RNA. RNA 30:124–135.

14. Masuda H, Tan Q, Awano N, Yamaguchi Y, Inouye M. 2012. A novel membrane-bound toxin for cell division, CptA (YgfX), inhibits polymerization of cytoskeleton proteins, FtsZ and MreB, in Escherichia coli. FEMS Microbiol Lett 328:174–181.

15. Tan Q, Awano N, Inouye M. 2011. YeeV is an Escherichia coli toxin that inhibits cell division by targeting the cytoskeleton proteins, FtsZ and MreB. Mol Microbiol 79:109– 118.

16. Wen Z, Wang P, Sun C, Guo Y, Wang X. 2017. Interaction of type IV toxin/antitoxin systems in cryptic prophages of Escherichia coli K-12. Toxins (Basel) 9:77.

17. Greer H. 1975. The kil gene of bacteriophage lambda. Virology 66:589–604.

18. Haeusser DP, Hoashi M, Weaver A, Brown N, Pan J, Sawitzke JA, Thomason LC, Court DL, Margolin W. 2014. The Kil peptide of bacteriophage λ blocks Escherichia coli cytokinesis via ZipA-dependent inhibition of FtsZ assembly. PLoS Genet 10:e1004217.

19. Hernández-Rocamora VM, Alfonso C, Margolin W, Zorrilla S, Rivas G. 2015. Evidence that bacteriophage λ Kil peptide inhibits bacterial cell division by disrupting FtsZ protofilaments and sequestering protein subunits. J Biol Chem 290:20325– 20335.

20. Geissler B, Elraheb D, Margolin W. 2003. A gain-of-function mutation in ftsA bypasses the requirement for the essential cell division gene zipA in Escherichia coli. Proc Natl Acad Sci 100:4197–4202.

21. Young KD. 2008. Why spherical Escherichia coli dies: the inside story. J Bacteriol 190:1497–1498.

22. Abramson J, Adler J, Dunger J, Evans R, Green T, Pritzel A, Ronneberger O, Willmore L, Ballard AJ, Bambrick J. 2024. Accurate structure prediction of biomolecular interactions with AlphaFold 3. Nature 1–3.

23. Erdős G, Dosztányi Z. 2020. Analyzing protein disorder with IUPred2A. Curr Protoc Bioinforma 70:e99.

24. Weber M, Burgos R, Yus E, Yang J, Lluch-Senar M, Serrano L. 2020. Impact of C-terminal amino acid composition on protein expression in bacteria. Mol Syst Biol 16:e9208.

25. Van Roey K, Uyar B, Weatheritt RJ, Dinkel H, Seiler M, Budd A, Gibson TJ, Davey NE. 2014. Short linear motifs: ubiquitous and functionally diverse protein interaction modules directing cell regulation. Chem Rev 114:6733–6778.

26. Lutkenhaus J. 1993. FtsZ ring in bacterial cytokinesis. Mol Microbiol 9:403–409.

27. McQuillen R, Xiao J. 2020. Insights into the structure, function, and dynamics of the bacterial cytokinetic FtsZ-ring. Annu Rev Biophys 49:309–341.

28. Govindarajan S, Nevo-Dinur K, Amster-Choder O. 2012. Compartmentalization and spatiotemporal organization of macromolecules in bacteria. FEMS Microbiol Rev.

29. Errington J. 2015. Bacterial morphogenesis and the enigmatic MreB helix. Nat Rev Microbiol 13:241–248.

30. Bendezú FO, Hale CA, Bernhardt TG, De Boer PAJ. 2009. RodZ (YfgA) is required for proper assembly of the MreB actin cytoskeleton and cell shape in E. coli. EMBO J 28:193–204.

31. Peters NT, Dinh T, Bernhardt TG. 2011. A fail-safe mechanism in the septal ring assembly pathway generated by the sequential recruitment of cell separation amidases and their activators. J Bacteriol 193:4973–4983.

32. Govindarajan S, Amster-Choder O. 2017. The bacterial Sec system is required for the organization and function of the MreB cytoskeleton. PLoS Genet 13.

33. Gueiros-Filho FJ, Losick R. 2002. A widely conserved bacterial cell division protein that promotes assembly of the tubulin-like protein FtsZ. Genes Dev 16:2544–2556.

34. Galli E, Gerdes K. 2010. Spatial resolution of two bacterial cell division proteins: ZapA recruits ZapB to the inner face of the Z-ring. Mol Microbiol 76:1514–1526.

35. Kruse T, Bork-Jensen J, Gerdes K. 2005. The morphogenetic MreBCD proteins of Escherichia coli form an essential membrane-bound complex. Mol Microbiol 55:78– 89.

36. Shiomi D, Sakai M, Niki H. 2008. Determination of bacterial rod shape by a novel cytoskeletal membrane protein. EMBO J 27:3081–3091.

37. Yakhnina AA, Gitai Z. 2012. The small protein MbiA interacts with MreB and modulates cell shape in Caulobacter crescentus. Mol Microbiol 85:1090–1104.

38. Fenton AK, Gerdes K. 2013. Direct interaction of FtsZ and MreB is required for septum synthesis and cell division in Escherichia coli. EMBO J 32:1953–1965.

39. Ursell TS, Nguyen J, Monds RD, Colavin A, Billings G, Ouzounov N, Gitai Z, Shaevitz JW, Huang KC. 2014. Rod-like bacterial shape is maintained by feedback between cell curvature and cytoskeletal localization. Proc Natl Acad Sci 111:E1025–E1034.

40. Govindarajan S, Borges A, Karambelkar S, Bondy-Denomy J. 2022. Distinct subcellular localization of a type i CRISPR complex and the Cas3 nuclease in bacteria. J Bacteriol 204:e00105–22.

41. Belitsky M, Avshalom H, Erental A, Yelin I, Kumar S, London N, Sperber M, Schueler-Furman O, Engelberg-Kulka H. 2011. The Escherichia coli extracellular death factor EDF induces the endoribonucleolytic activities of the toxins MazF and ChpBK. Mol Cell 41:625–635.

42. Edgar RC. 2004. MUSCLE: multiple sequence alignment with high accuracy and high throughput. Nucleic Acids Res 32:1792–1797.

43. Kumar S, Tamura K, Nei M. 2004. MEGA3: Integrated software for Molecular Evolutionary Genetics Analysis and sequence alignment. Brief Bioinform 5:150–63.

44. Stamatakis A. 2014. RAxML version 8: a tool for phylogenetic analysis and post-analysis of large phylogenies. Bioinformatics 30:1312–1313.

45. Letunic I, Bork P. 2007. Interactive Tree Of Life (iTOL): an online tool for phylogenetic tree display and annotation. Bioinformatics 23:127–8.

